# Modeling the effects of motivation on choice and learning in the basal ganglia

**DOI:** 10.1101/797860

**Authors:** Maaike M.H. van Swieten, Rafal Bogacz

## Abstract

Decision making relies on adequately evaluating the consequences of actions on the basis of past experience and the current physiological state. A key role in this process is played by the basal ganglia, where neural activity and plasticity are modulated by dopaminergic input from the midbrain. Internal physiological factors, such as hunger, scale signals encoded by dopaminergic neurons and thus they alter the motivation for taking actions and learning. However, to our knowledge, no formal mathematical formulation exists for how a physiological state affects learning and action selection in the basal ganglia. We developed a framework for modelling the effect of motivation on choice and learning. The framework defines the motivation to obtain a particular resource as the difference between the desired and the current level of this resource, and proposes how the utility of reinforcements depends on the motivation. To account for dopaminergic activity previously recorded at different physiological states, the paper argues that the prediction error encoded in the dopaminergic activity needs to be redefined as the difference between utility and expected utility, which depends on both the objective reinforcement and the motivation. We also demonstrate a possible mechanism by which the evaluation and learning of utility of actions can be implemented in the basal ganglia network. The presented theory brings together models of learning in the basal ganglia with the incentive salience theory in a single simple framework, and it provides a mechanistic insight into how decision processes and learning in the basal ganglia are modulated by the motivation. Moreover, this theory is also consistent with data on neural underpinnings of overeating and obesity, and makes further experimental predictions.

**Author summary:** Behaviour is made of decisions that are based on the evaluation of costs and benefits of potential actions in a given situation. Actions are usually generated in response to reinforcement cues which are potent triggers of desires that can range from normal appetites to compulsive addictions. However, learned cues are not constant in their motivating power. Food cues are more potent when you are hungry than when you have just finished a meal. These changes in cue-triggered desire produced by a change in biological state present a challenge to many current computational models of motivation and learning. Here, we demonstrate concrete examples of how motivation can instantly modulate reinforcement values and actions; we propose an overarching framework of learning and action selection based on maintaining the physiological balance to better capture the dynamic interaction between learning and physiology that controls the incentive salience mechanism of motivation for reinforcements. These models provide a unified account of state-dependent learning of the incentive value of actions and selecting actions according to the learned positive and negative consequences of those actions and with respect to the physiological state. We propose a biological implementation of how these processes are controlled by an area in the brain called the basal ganglia, which is associated with error-driven learning.

## Introduction

Successful interactions with the environment rely on using previous experience to predict the value of outcomes or consequences of available actions. Human and animal studies have strongly implicated the neurotransmitter dopamine in these learning processes [1–8], in addition to its roles in shaping behaviour, including motivation [9], vigour [10] and behavioural activation [11, 12].

Dopamine seems to have two distinct effects on the networks it modulates. First, it facilitates learning by triggering synaptic plasticity [13]. Such dopaminergic teaching signal is thought to encode a reward prediction error (RPE), which is defined as a difference between a reinforcement and the expected reinforcement [1]. The overall value of a reinforcement that is available at a given moment depends on the potential positive and negative consequences associated with obtaining it. These consequences can be influenced by internal and external factors, such as the physiology of the subject and the reinforcement’s availability, respectively. Information about the external factors is indeed encoded in the dopaminergic responses which are shown to scale with the magnitude and the probability of a received reinforcement [14, 15], but also with the delay and effort related costs associated with a reinforcement [16, 17]. Second, the level of dopamine controls the activation of the basal ganglia network by modulating excitability of neurons [18, 19]. Although dopamine is a critical modulator of both learning and activation, it is unclear how it is able to do both given that these processes are conceptually, computationally and behaviourally distinct. For a long time, our understanding was that tonic (sustained) levels of dopamine encode an activation signal and phasic (transient) responses convey a teaching signal (i.e. prediction error) [10]. However, recent studies have shown that this distinction is not as clear as we thought [20, 21] and that other mechanisms may exist, which allow striatal neurons to correctly decode the two signals from dopaminergic activity [22]. In this paper, we do not investigate mechanisms by which these different signals can be accessed, but we assume that striatal neurons can read out both activation and teaching signals encoded by dopaminergic neurons.

In addition to external factors explained above, internal physiological factors, such as hunger, can also alter the reinforcement value of an action and drive decision making based on the usefulness of that action and the outcome at that given time. For example, searching for food when hungry is more valuable than when sated and actions have to be evaluated accordingly. In other words, the current physiological state affects the motivation to obtain a particular resource. The physiological state has indeed been observed to modulate dopamine levels and dopamine responses encoding reward prediction error [23–25], thus it is likely that the physiological state influences both the activation and teaching signals carried by dopamine. Strikingly, the physiological state can sometimes even reverse the value of a reinforcement (e.g. salt) from being rewarding in a depleted state to aversive in a sated state [26]. Moreover, the physiological state during learning may affect subsequent choices, for example, animals may still have a preference for actions that were associated with hunger even when they are sated [27]. Recently, it has been proposed how physiological state can be introduced into reinforcement learning theory to refine the definition of a reinforcement [28]. However, despite the importance of the physiological state for describing behaviour and dopaminergic activity, we are not aware of theoretical work that integrates the physiological state into a theory of dopaminergic responses.

Another important line of work describing subjective preferences is the utility theory. It is based on the assumption that people can consistently rank their choices depending upon their preferences. The utility theory has been used extensively in economics [29], and it has been shown that dopaminergic responses depend on the subjective utility of the obtained reward magnitude, rather than its objective magnitude [30]. As described above, there is a need to extend the general utility function with a motivational component that describes the bias in the evaluation of positive and negative consequences of decisions as a result of changes in the physiological state of a subject. Evidence for this bias comes from devaluation studies in which reinforcements are specifically devalued by pre-feeding or taste aversion. The concept of state-dependent valuation has been studied in various contexts [24, 31, 32] and in different species, including starlings [33, 34], locusts [35] and fish [27]. These studies suggest that the utility of outcomes depends on both the (learned) reinforcement value and the physiological state. One of the earliest attempts to capture this relationship between incentive value and internal motivational state is the incentive salience theory [12].

In this paper we aim to provide an explanation for the above effects of physiological state on behaviour and dopaminergic activity with a simple framework that combines incentive learning theory [36, 37] with models of learning in the basal ganglia. By integrating key concepts from these theories we define a utility function for actions that can be modulated by internal and external factors. In our framework, the utility is defined as the change in the desirability of physiological state resulting from taking an action and obtaining a reinforcement. Following previous theoretic work [28], the motivation for a particular resource is defined as the difference between the desired and the current level of this resource.

In the proposed framework, motivation affects both teaching and activation signals encoded by dopaminergic neurons. Relying on experimental data, we argue that the dopaminergic teaching signal encodes the difference between utility and expected utility, which depends on motivation. Moreover, we propose how motivation can influence the dopaminergic activation signal to appropriately drive action selection behaviour. We also highlight that the resulting consequences of an action can be positive or negative depending on how far the current and new physiological state are from the desired state. Building on existing theories we illustrate how the neurons in the striatum could learn these consequences through plasticity rules. Finally, we use the resulting models to explain experimental data. Together, this paper discusses a modelling framework that describes how the internal physiological state affects learning and action selection in the basal ganglia and provides novel interpretations of existing experimental data. To provide a rationale for our framework the remainder of the introduction reviews the data on effects of physiological state on dopaminergic teaching signal.

### Effects of motivation on dopaminergic responses

We first review a classical reinforcement learning theory and then discuss data which challenges it. As postulated by reinforcement learning theories, expectations of outcomes are updated on the basis of experiences. This updating process may be guided by prediction errors, which are computed by subtracting the received reinforcement (*r*) from the cached reinforcement expectation (*V_t_*). In classical conditioning and after extensive training, the dopamine response to the conditioned stimulus (CS) is observed to reflect the expected future reinforcement, whereas the response to the unconditioned stimulus (US) represents the difference between the obtained reinforcement and the expectation [1]. To account for these responses, the reward prediction error in a temporal difference model (*δ_TD_*) is classically defined as [1]:

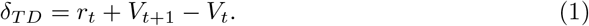

The above equation defines the prediction error as the difference between total reinforcement (including both reinforcement actually received *r_t_* and reinforcement expected in the future *V*_*t*+1_) and the expected reinforcement (*V_t_*).

We now review how the above equation captures the dopamine responses at the time of the CS and the US, which change over the course of learning. At the start of learning, the animal has not formed any expectation yet, which means that at the time of the CS, *V_t_* is 0. Given that no reinforcement is provided at the time of CS presentation, *r_t_* is also 0. Thus, the prediction error at the time of the CS is equal to the expected value of the reinforcement (*δ_TD_* = *V*_*t*+1_). The response to the CS is zero in naive animals. By contrast, the response to the CS in fully trained animals reflects the expected upcoming reinforcement, as extensive training allowed animals to update their expectations to predict upcoming reinforcements better. At the time of the US, no future reinforcements are expected so *V*_*t*+1_ is 0, thus the reward prediction error at the time of US is equal to *δ_TD_* = *r_t_* − *V_t_*. Unpredicted rewards (i.e. positive reinforcements) evoke positive prediction errors, while predicted reinforcements do not. Thus this definition of prediction error captures observed patterns of dopaminergic responses; where naive animals, which are unable to predict reinforcements, show large positive responses at the time of the US, fully trained animals that can predict reinforcements perfectly, show no response at all.

Recently, the above definition of reward prediction error has been experimentally challenged by Cone et al. [25]. They show that the internal state of an animal modulates the teaching signals encoded by dopamine neurons in the midbrain after conditioning (Fig 1). In this study, animals were trained and tested in either a sodium depleted or sodium balanced state. The dopaminergic responses predicted by the classical reinforcement learning theory shown by Schultz et al. [1] were only observed in animals which were both trained and tested in the depleted state. In all other conditions the dopaminergic responses followed different patterns. When animals were trained in the balanced state but tested in the depleted state, increased dopaminergic responses to the US (i.e. salt infusion) rather than the CS were observed, which is similar to dopaminergic responses observed in untrained animals in the study of Schultz et al. [1], suggesting that learning did not occur in the balanced state. When animals were trained and tested in the balanced state, there was no dopaminergic response to either CS or US. Interestingly, the same pattern was observed in animals trained in the depleted state and tested in the balanced state, suggesting that the learned values are modulated. In the Results section below we will demonstrate how this pattern of activity can be captured by appropriately modifying a definition of prediction error.

**Fig 1.**
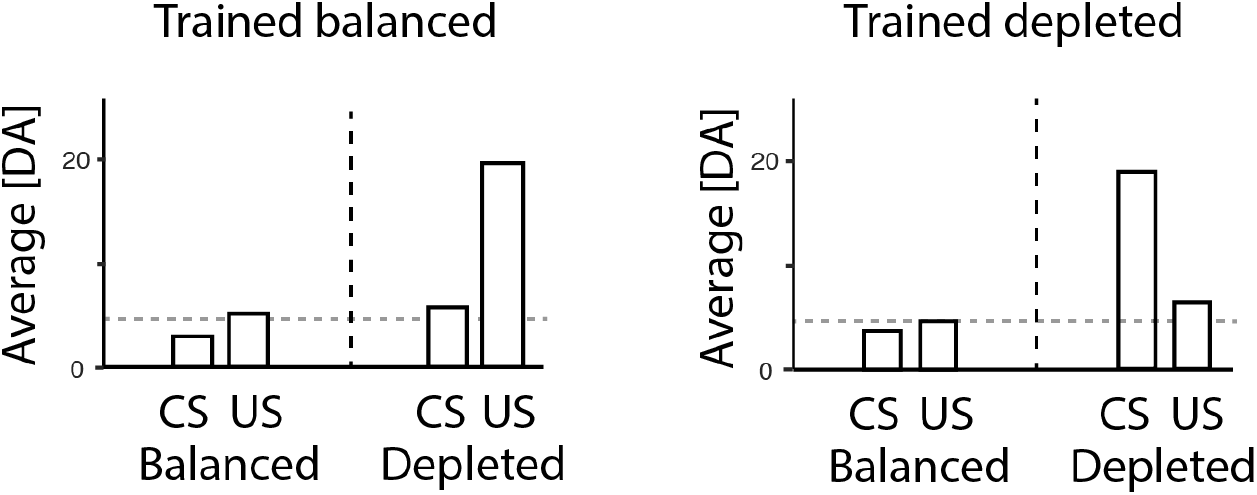
Experimental data by Cone et al. [25]. The two graphs within the figure correspond to dopaminergic responses in animals trained in a balanced and depleted state, respectively, re-plotted from figures 2 and 4 in the paper by Cone et al., (2016). Within each graph, the left and right halves show the responses of animals tested in balanced and depleted states, respectively. The horizontal dashed lines indicate baseline levels.

## Results

In this section, we present our framework and its possible implementation in the basal ganglia circuit, and illustrate how it can account for the effects of motivation on neural activity and behaviour.

### Normative theory of state-dependent utility

The utility and consequences of actions are dependent on the usefulness of the reinforcement (*r*) with respect to the current state. To maintain a physiological balance the distance between the current state *S* and the desired state *S*\* has to be minimised. We assume that the desirability function of a physiological state has a concave, quadratic shape (Fig 2A), because it is more important to act when you are in a very low physiological state, compared to when in a near optimal state. Thus, we define a desirability of a state in the following way (a constant of 1/2 is added for mathematical convenience, as it will cancel in subsequent derivations):

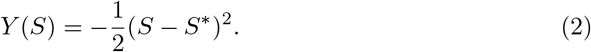

**Fig 2.**
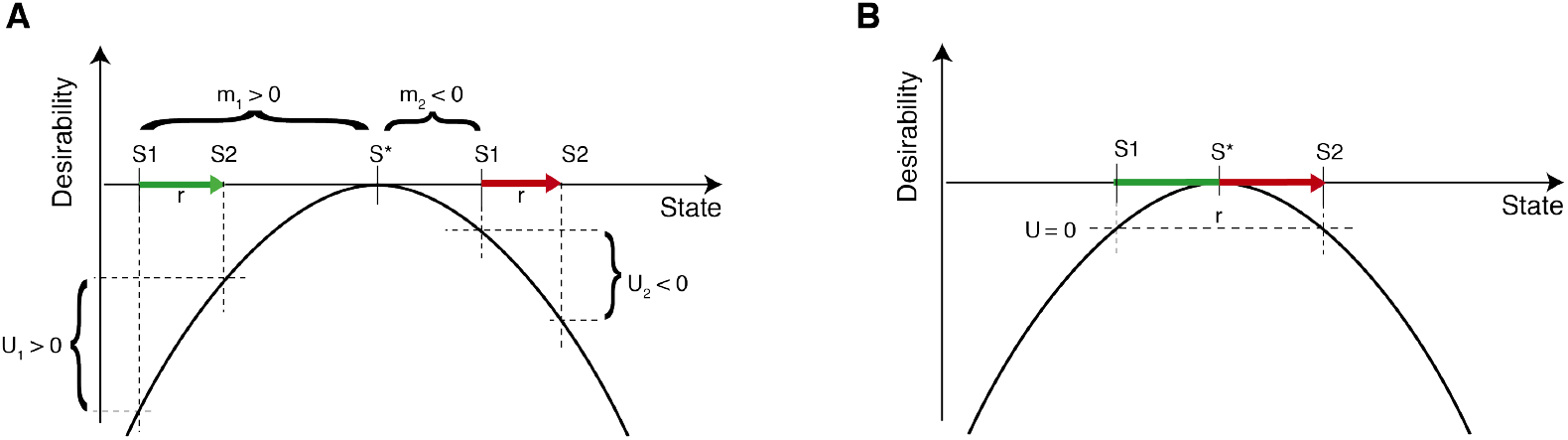
Dependence of utility of an action on the reinforcement and physiological state. A) Same reinforcement may have positive or negative utility depending on the physiological state. B) A large reinforcement may have no utility even if the animal is initially in a depleted state. U = utility, m = motivation. S* = the desired state and S1 = state before action, S2 = state after action. Arrow length indicates the size of the reinforcement (*r*). Changes in state resulting in increase and decrease of desirability are indicated with green and red arrows, respectively.

We define the utility *U* of an action as a change in desirability of the physiological state resulting from taking that action. Fig. 2A illustrates that the utility of an action depends on both the obtained reinforcement *r* for that action and the motivation *m*, which is defined as the difference between the desired and the current physiological state:

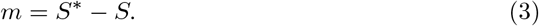

According to the above definition, the same reinforcement could yield a positive or negative utility of an action, depending on whether the difference between the current physiological state and the desired state is positive or negative (Fig. 2A). This parallels an observation that nutrients such as salt may be appetitive or aversive depending on the level of an animal’s reserves [26]. Although not discussed in this paper, please note that this definition of the utility also can be extended to the utility of an external state, such as a particular location in space. The utility of such an external state can be defined as a utility of the best available action in this state.

Before presenting an exact expression, it is useful to consider a simple approximate expression for the utility. Such approximation can be obtained through a first order Taylor expansion of Eq. (2):

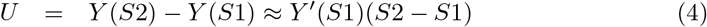

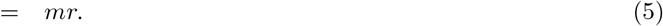

This approximation of the utility clearly shows that the utility of an action, defined as the change in physiological state, depends on both the motivation *m* and the reinforcement *r*, where *r* = *S*2 − *S*1.

In order to select actions on the basis of their utility, animals needs to maintain an estimate of the utility *Û* of an action. There are several ways such an estimate can be learned. Here we discuss a particular learning algorithm, which results in prediction errors that resemble those observed by Cone et al. [25]. This learning algorithm assumes that animals minimise the absolute error in the prediction of the utility of the chosen action. We can therefore define this prediction error as:

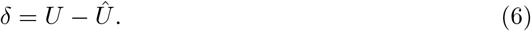

The above expression for the prediction error (Eq. (6)) provides a general definition of the prediction error as the difference between the observed and expected utility. In this paper we claim that this expression better describes the dopaminergic teaching signal observed in experimental data, which we will demonstrate in more detail in the next section.

Assuming that the animal’s estimate of expected reinforcement is encoded in a parameter *V*, the animal’s estimate of the utility is *Û* = *mV*. Combining Eq. (5) with Eq. (6), we obtain the following expression for the reward prediction error:

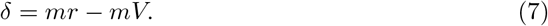

To predict upcoming reinforcement better, the absolute prediction error has to be minimised. We can define an objective function that will be maximised:

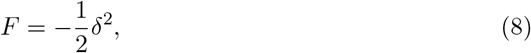

In order to maximise this objective function, the estimate of the expected reinforcement, *V*, is updated proportionally to the prediction error:

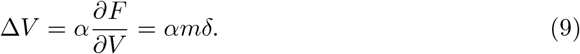

### Simulating state-dependent dopaminergic responses

This section serves to illustrate that the pattern of dopaminergic activity seen in the study by Cone et al. [25] is not consistent with the classical theory and can be better explained with a state-dependent utility as described above. We first simulated the classical model in which reward prediction error is described in Eq. (1). In the simulation, the CS was presented at time step 1, while the US was presented at time step 2. The model was learning a single parameter *V* estimating the expected reinforcement on the time step following the CS. Thus, on each trial the prediction error to the CS was equal to *δ_TD_* = *V*, while the prediction error to the US was equal to *δ_TD_* = *r* − *V*. The value estimate was updated proportionally to the prediction error, i.e. Δ*V* = *α*(*r* − *V*), where *α* is a learning rate parameter. On every trial, the model received a reinforcement *r* = 0.5. Once training was completed, expected values were fixed to the values they converged to during training, and testing occurred without allowing the model to update the beliefs.

Since the classical reward prediction error does not depend on the physiological state, each experimental condition was simulated in exactly the same way, and dopaminergic teaching signals, predicted by the classical theory, are identical in all conditions (Fig 3A). An estimate of the expected value of the reinforcement is reflected by a response to the CS and the response to the US is close to zero as the reinforcement received is fully predicted.

**Fig 3.**
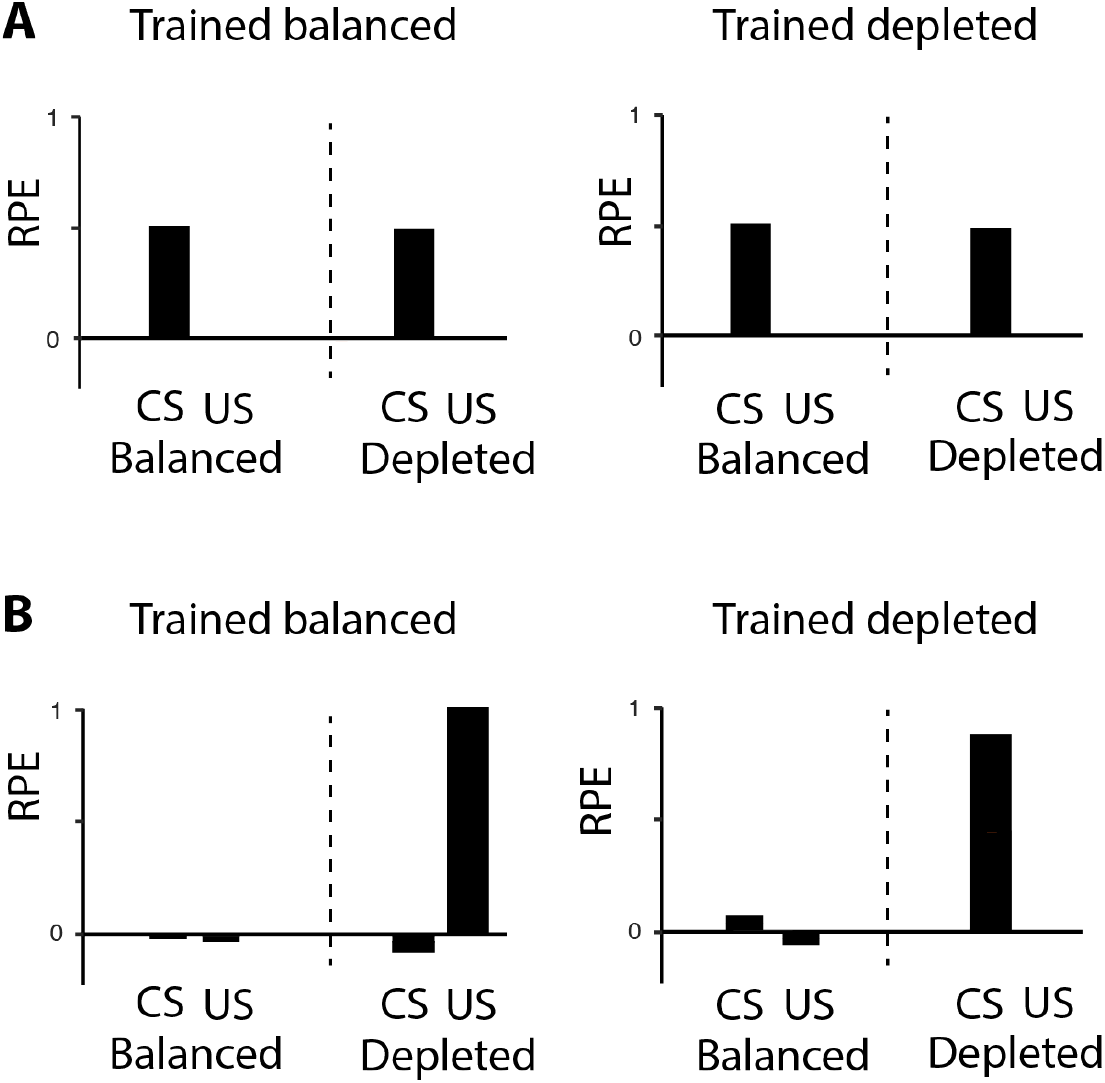
Simulated data by Cone et al. [25] with different reward prediction errors. A) Simulation with typical reward prediction error using Eq. (1). B) Simulation with scaled utility and scaled expectation using Eq. (7). Within each graph, the left and right halves show the reward prediction error (RPE) of simulated animals tested in balanced and depleted states, respectively. CS = conditioned stimulus, US = unconditioned stimulus. Each simulation consisted of 50 trials and was repeated 5 times, similar to the number of animals in each group in the study by Cone et al. (2016). The learning rate for this experiment was set to *α* = 0.1. Error bars are equal to zero as there is no noise added to the simulation.

The simulations employing the state-dependent prediction error (Eq. (7)) as defined in the previous section followed the same protocol as in the classical case. During training, at the time of US the reinforcement estimate was updated proportionally to the prediction error, Δ*V* = *αδ*. This update is similar to that in Eq. (9), but for simplicity was not scaled by m. Adding this scaling factor does not qualitatively change the resulting pattern of the prediction error as m was a positive constant in all the simulations.

During testing, the values were no longer modified, and the dopaminergic teaching signal at the time of the US was computed from Eq. (7), while the value at the time of the CS was taken as *mV*. The parameter describing motivation was set to *m* = 0.2 for a state close to balanced and *m* = 2 for a depleted state.

In simulated animals that are trained in the near-balanced state little learning is triggered and the response to the CS is close to zero (Fig 3B). However, when these simulated animals are then tested in the depleted state, the scaled utility is greater than zero and consequently evokes a positive reward prediction error. In contrast, simulated animals trained in the depleted state learn the estimate of the expected value of the reinforcement. There is an increase in the dopaminergic teaching signal in these simulated animals at the time of the CS since the expected value is transferred to the CS. When these simulated animals are tested in the near-balanced state, with a motivation close to zero, a very small reward prediction error is evoked, because both the reinforcement and expected value are scaled by a number close to zero.

In line with the theory in the previous section in which we formally defined both the utility and motivation, the above simulations shows that in order to account for the experimental data by Cone et al., (2016), the prediction error needs to be redefined as a difference between the utility of a reinforcement and the expected utility of that reinforcement, which depends on both the objective reinforcement magnitude and the motivation.

### Accounting for positive and negative consequences of actions

Let us now reconsider how the dependence of utility on motivation may be expressed more accurately. Since Eq. (4) comes from Taylor expansion, it only provides a close approximation if r is small. This approximation may fail when the reinforcement is greater than the distance to the optimum. In the example in Fig 2B, if we use a linear approximation with a positive motivation, the utility is approximated as greater than zero, even though the actual utility is not as this action will exceed the desired state. Eq. (4) also suggests that any action with *r* > 0 will have positive utility if *m* > 0, regardless of possible negative consequences (i.e. reaching a new state further away from the desired state). Moreover, if the distance of the current state to the desired state is equal to the distance of the new state to the desired state, the utility of an action would be zero (Fig. 2B). Using Eq. (4) it is impossible to capture these effects and account for both positive and negative consequences of this action.

One classical example in which the utility of an action switches sign depending on the proximity to the desired physiological state is salt appetite. When animals are depleted of sodium, salt consumption is rewarding. However, when animals are physiologically balanced, salt consumption is extremely aversive [26]. To avoid using multiple equations to explain the switch from positive to negative utilities and vice versa [37], we need to formulate an equation that can account for negative consequences of actions when the *m* ≈ 0 or *m* < 0 and is able to account for positive consequences when the motivation changes, i.e. *m* > 0.

Therefore we use a second order Taylor expansion which gives an exact expression for the utility:

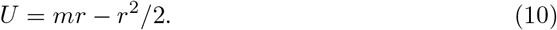

In the above equation *mr* could be seen as the positive and *r*^2^/2 as the negative consequences of the action, respectively. The first term plays a greater role when deprived and promotes taking actions, whereas the second term plays a greater role when balanced and discouraged taking actions.

During action selection it is imperative to choose actions to maximise future utility. For competing actions, the utility of all available actions needs to be computed. The action with the highest utility is most beneficial to select, but this action should only chosen when its utility is positive. If the utility of all actions is negative, no actions should be taken. From a fitness point of view, not making an action is more advantageous than incurring a high cost.

In the next section we will elaborate on how Eq. (10) can be evaluated in the basal ganglia and provide an example of a biologically plausible implementation. For simplicity we will only consider a single physiological dimension (e.g. nutrient reserve), but we recognise that the theory needs to be extended in the future to multiple dimensions (e.g. water reserve, fatigue) which an animal needs to optimise. Furthermore, taking an action aimed to restore one dimension (e.g. nutrient reserve) may also include negative consequences that are independent of the considered dimension (e.g. fatigue). We will elaborate on these issues in the Discussion.

### Neural implementation

In the previous sections we discussed how the utility of actions or stimuli change in a state-dependent manner. In this section we will focus on the neural implementation of these concepts. More specifically, we will address how the utility of previously chosen actions can be computed in the basal ganglia and how this circuit could learn the utility of actions.

#### Evaluation of utility in the basal ganglia circuit

The basal ganglia is a group of subcortical nuclei that play a key role in action selection and reinforcement learning. It is organised into two main pathways shown schematically in Fig 4. The Go or direct pathway is associated with the initiation of movements, while the Nogo or indirect pathway is associated with the inhibition of movements [38]. These two pathways include two separate populations of striatal neurons expressing different dopaminergic receptors [39]. The striatal Go neurons mainly express D1 receptors which are excited by dopamine, while the striatal Nogo neurons mainly express D2 receptors which are inhibited by dopamine [40]. Thus, dopaminergic activation signal controls the competition between these two pathways during action selection and promotes action initiation over inhibition.

**Fig 4.**
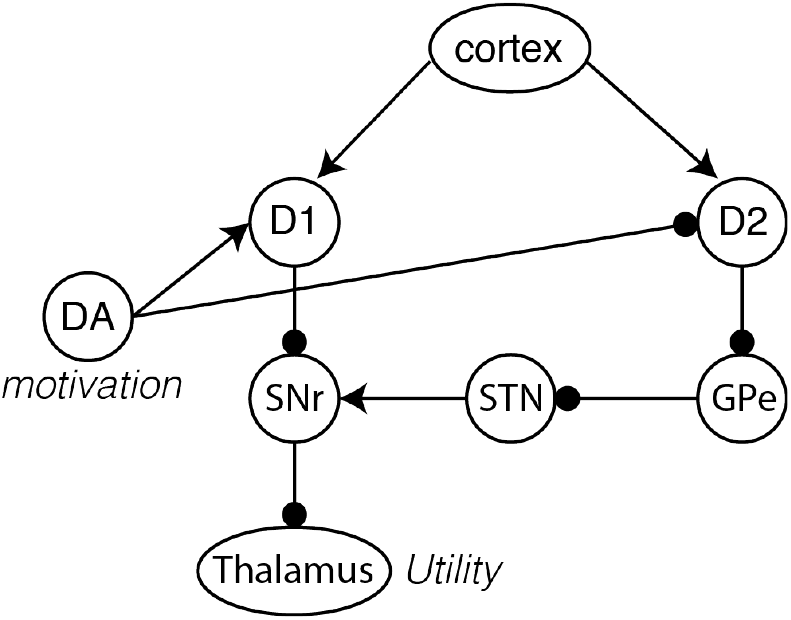
Schematic of utility computation in the basal ganglia network. Dopaminergic activation signal encodes the motivation. The thalamic activity represents the utility of actions. Arrows and lines with circles denote excitatory and inhibitory connections, respectively. DA = dopamine, D1 = Dopamine receptor 1 medium spiny neurons, D2 = Dopamine receptor 2 medium spiny neurons, SNr = Substantia Nigra Pars Reticulata, STN = Subthalamic Nucleus, GPe = Globus pallidus external segment.

Given the architecture of the basal ganglia, we hypothesise that this circuitry is well suited for the computation of the utility of actions in decision making. This utility could be encoded at the final processing stage of this network, i.e. the thalamus. In particular, we suggest that the Go neurons will mostly determine thalamic activity when the utility is positive, while, the Nogo neurons when the utility is negative, and the dopaminergic activation signal can appropriately control the relative influence of Go and Nogo neurons, because it encodes motivation. There are various ways to describe how the utility is represented in the basal ganglia and how the basal ganglia output can drive action selection. In this paper, we show one possibility that should only be treated as a proof of principle.

In line with earlier studies, we assume that synaptic strengths of Go and Nogo neurons encode positive and negative consequences of actions, respectively [41–43]. Accordingly, in the presented model the Go and Nogo neurons represent the estimates for the two terms in Eq. (10), namely, Go neurons produce activity proportional to *mr* while Nogo neurons produce activity proportional to *r*^2^/2.

We refer to the output of the basal ganglia as the thalamic activity, denoted by *T*. *T* depends on the cortico-striatal weights of Go neurons (*G*) and Nogo neurons (*N*), and the dopaminergic activation signal denoted by *D*. The striatal weights of Go neurons have an overall positive effect on the thalamic activity as the projection from the Go neurons to the thalamus involves a double inhibitory connection. In contrast, the inhibitory effect of Nogo neurons on the thalamic activity result in a negative contribution to the thalamic activity. We assume that the dopaminergic activation signal increases the gain of Go neurons, based on the observation of an increased slope of firing-input relationship of neurons expressing D1 receptors in the presence of dopamine [18]. In contrast, we assume that the dopaminergic activation signal reduces the gain of Nogo neurons, as their firing-input relationship has decreased slope in the presence of dopamine [19].

Although admittedly more complex, we can capture the signs of the influences of the dopaminergic activation signal, Go and Nogo neurons in a linear approximation [43]:

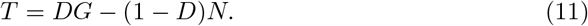

In the above equation, the contribution of Go neurons to the thalamic activity is described by the first term *DG*, reflecting facilitatory effect of dopamine on Go neurons. The inhibitory connection of Nogo neurons to the thalamic activity results in a negative contribution to the thalamic activity and is described by the second term −(1 − *D*)*N*. We assume that *D* ∈ [0, 1], meaning that a value of *D* = 0.5 corresponds to a baseline level of dopaminergic activation signal for which both striatal populations equally contribute to the thalamic activity.

We now show that the thalamic activity defined in Eq. (11) is proportional to the utility of an action if *G* and *N* are fully learned and therefore provide correct estimates of the positive and negative terms in utility equation (Eq. (10)), respectively (*G* = *r* and *N* = *r*^2^/2). Then, we can rewrite Eq. (11) as:

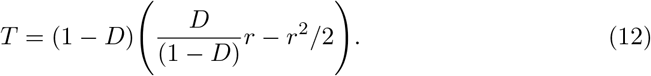

Comparing this to Eq. (10), we observe that the thalamic activity is proportional to the utility (*T* = (1 − *D*)*U*) when the motivation is encoded by dopaminergic activation signal:

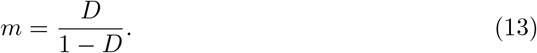

We can rewrite Eq. (13) in the following way to express the level of dopamine for a given motivation:

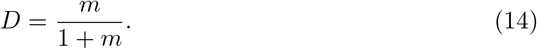

In summary, when the striatal weights encode the positive and negative consequences and the dopaminergic activation signal is described by Eq. (14), then the thalamic activity is proportional to the utility.

Let us now consider how this utility can be used to guide action selection. Computational models of action selection typically assume that all basal ganglia nuclei and thalamus include neurons selective for different actions [44]. Therefore, the activity of thalamic neurons selective for specific actions can be determined on the basis of their individual positive and negative consequences and the common dopaminergic activation signal. Given that the proportionality coefficient (1 − *D*) in Eq. (12) is the same for all actions, the utilities of different actions represented by thalamic activity are scaled by the same constant. This means that the most active thalamic neurons are the ones selective for the action with the highest utility, and hence this action may be chosen through competition. Furthermore, if we assume that actions are only selected when thalamic activity is above a threshold, then no action will be selected if all actions have insufficient utility. The utility of actions has to be sufficiently high to increase neural firing in the thalamus above the threshold and trigger action initiation.

#### Models of learning

In the previous section, we showed that the basal ganglia network can estimate the utility once the striatal weights have acquired the appropriate values. In this section we address the question of how these values are learned. Earlier, we proposed a general framework for describing learning process assuming that the brain minimises a prediction error during this process and we redefined the prediction error as the difference between utility and expected utility. In the previous section we described a model in which the thalamic activity encodes a scaled version of the estimated utility (*Û* = *T*/(1 − *D*) (see Eq. 12 and 13). This estimate of the utility can be substituted into Eq. (6) giving the following the state-dependent reward prediction error:

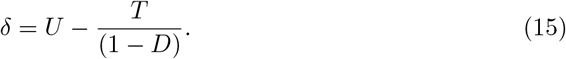

In this RPE, *U* is the utility of an action (Eq. (10)), and *T/*(1 − *D*) is the expected utility of an action. In line with the general definition of Eq. (6), the above equation shows that the RPE depends on a reinforcement and the expected reinforcement in a state-dependent manner, which is here instantiated in a specific model for estimating utility. Please note that at baseline levels of the dopaminergic activation signal (*D* = 0.5), the above expression for prediction error reduces to *δ* = *U* − (*G* − *N*) and such prediction error has been used previously [42].

We will now describe two models for learning the synaptic weights of *G* and *N*. The first model is a normative model, developed for the purpose of this learning, while the second model corresponds to a previously proposed model of striatal plasticity, and it provides a more biologically realistic approximation of the first model.

##### Gradient model

The first model we use to describe learning of synaptic weights under changing conditions, directly minimises the error in prediction of the utility of action. It changes the weights proportionally to the gradient of the objective function:

Δ*G* = *α∂F*/*∂G* and Δ*N* = *α∂F*/*∂N*, respectively. For the prediction error described in Eq. (15), this gives us the following learning rules for *G* and *N*:

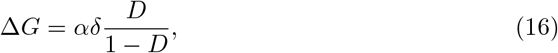

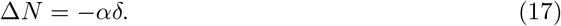

Synaptic weights of Go and Nogo neurons are updated using the dopaminergic teaching signal scaled by the learning rate constant *α*. The update rule for Go weights has an additional term involving the dopaminergic activation signal encoding the motivation as described in Eq. (13). Only the update rule for *G*, but not for *N*, includes scaling by motivation, because in the definition of utility of Eq. (10), the motivational level only scales the positive consequences of an action and not the negative.

##### Payoff-cost model

The second model has been previously proposed to describe how Go and Nogo neurons learn about payoffs and costs of actions. It has been shown to account for a variety of data ranging from properties of dopaminergic receptors on different striatal neurons to changes in risk preference when dopamine levels are low or high [42]. We expected this model to provide an approximation for the gradient model because it has been shown to be able to extract positive and negative consequences of actions. More specifically, if reinforcement takes positive value *r_p_* half of the times and negative value −*r_n_* the other half of times, then the Go weights converge to *G* = *r_p_* and Nogo weights to *N* = *r_n_*, for certain parameters [43]. Therefore, we expected this learning model to be able to extract positive and negative terms of the utility in Eq. (10) if motivation could vary between trials, so the positive term dominates utility on some trials while the negative term on other trials.

In our simulations we used the same update rules as previously described [42, 43], but we use a state-dependent prediction error (Eq. (15)) to account for decision making under different physiological states.

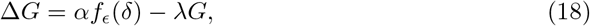

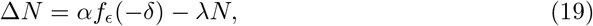

where

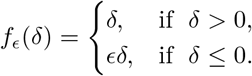

The update rules in the above equations consist of two terms. The first term is the change depending on the dopaminergic teaching signal scaled by a learning rate constant *α*. It increases the weights of Go neurons when *δ* > 0, and slightly decreases when *δ* < 0, so that changes in the Go weights mostly depend on positive prediction errors. The constant *ϵ* controls the magnitude by which the weights are decreased. Nogo weights will be updated in a analogous way, but these changes mostly depend on negative prediction errors. The second term in the update rules is a decay term, scaled by a decay rate constant *λ*. This term is necessary to ensure that the synaptic weights stop growing when they are sufficiently high and allows weights to adapt more rapidly when conditions change. In case an updated weight becomes negative, it is set to zero.

#### Simulations of learning

In this section, we investigate under what conditions the learning rules described above can yield synaptic weights of Go and Nogo neurons that allow for the estimation of utility. Recall that the network will correctly estimate the utility, if *G* = *r* and *N* = *r*^2^/2.

In the simulations we make the following assumptions: 1) The simulated animal knows its motivational level *m*, which influences both dopaminergic signals accordingly (Eqs. (14) and (15)). 2) The simulated animal computes the utility of obtained reinforcement as a change in the desirability of the physiological state. As described above, the desirability depends on the objective value of the reinforcement *r* and the current motivational state *m* according to Eq. (10), which was used to compute the reward prediction error according to Eq. (15).

We simulated scenarios in which the simulated animal repeatedly chooses a single action and experiences a particular reinforcement *r* under different levels of motivation *m* ∈ {*m*_low_, *m*_baseline_, *m*_high_}. Note that the *m*_low_ = 0 correspond to a dopaminergic activation signal of *D* = 0, *m*_baseline_ = 1 gives a dopaminergic activation signal of *D* = 0.5, which means that Go and Nogo neurons are equally weighted, and *m*_high_ = 2 corresponds to a dopaminergic activation signal above baseline levels.

We first simulated a condition in which the motivation changed on each trial, and took a randomly chosen value from a set {*m*_low_, *m*_baseline_, *m*_high_} (Fig 5A). The gradient model was able to learn the desired values of Go and Nogo weights. In particular, Go weights converged to *r*, while Nogo weights converged to *r*^2^/2, which allowed the network to correctly estimate the utility. Although the subjective reinforcing value changed as a function of physiological state, the model was able to learn the actual reinforcement of an action. Encoding of such objective estimates allows the agent to dynamically modulate behaviour based on metabolic reserves.

**Fig 5.**
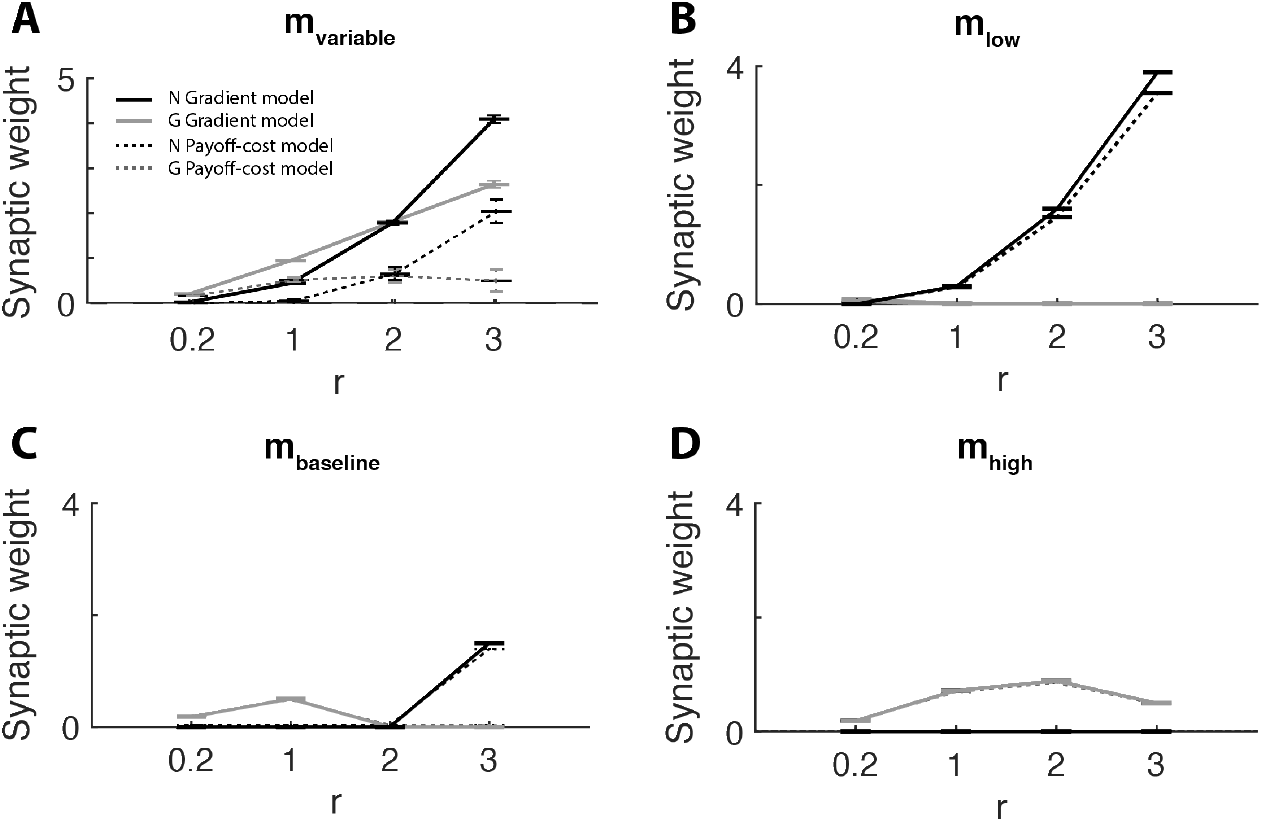
Learning Go and Nogo weights for different reinforcements and different levels of motivation. Performance of models under variable (A), low (B), baseline (C), high (D) levels motivation. Simulations were performed using the state-dependent prediction error (Eq. (15)). Solid lines show simulations of the gradient model using the plasticity rules described in Eq. (16) and (17). Dashed lines show simulations of the payoff-cost model using the plasticity rules described in Eq. (18) and (19). Black lines correspond to Nogo neurons and grey lines to Go neurons. Each simulation had 150 trials and was repeated 100 times. All synaptic weights were initialised at zero. The parameters used in the simulations were *α* = 0.1, *ϵ* = 0.8 and *λ* = 0.01. These parameters allow the model to converge to positive and negative consequences at baseline motivational state [43].

In contrast, the payoff-cost model converged to lower weights than desired. Although it learned the synaptic weights based on the state-dependent prediction error, the weight decay present in the model resulted in a lower asymptotic value.

To test robustness of the learning rules and because the motivational state is fixed during the experimental paradigms simulated in this paper, we also simulated conditions in which the motivation was kept constant (Fig 5B-D). In these cases both leaning rules converged to very similar values of synaptic weights: low levels of motivation emphasised negative consequences and therefore facilitated Nogo learning (Fig 5B), while high levels of motivation emphasised positive consequences and therefore facilitated Go learning (Fig 5D).

In summary, the simulations indicate that for the models to learn appropriate values of synaptic weights, the reinforcements need to be experienced under varying levels of motivation. In this case, the gradient model provides a precise estimation, while the payoff-cost model provides an approximation of the utility. In cases when the motivational state is fixed during training, both models learn very similar values of the weights.

#### The basal ganglia architecture allows for efficient learning

In the previous sections we presented and analysed models of how utilities can be computed and learned in the basal ganglia network. One could ask, why would the brain employ such complicated mechanisms if a simple model could give you the same results? In particular, one could consider a standard Q-learning model, in which the state is augmented by motivation. Such model would also be able to learn to estimate the utility. However, such a model does not incorporate any prior knowledge about the form of the utility function and its dependence on motivation. By contrast, the model grounded in basal ganglia architecture, assumes a particular form of the utility function to be learned. In machine learning, such prior assumptions are known as ‘inductive bias’, and they facilitate learning [45].

We now illustrate that thanks to the correct inductive bias, the gradient model learns to estimate the utility faster than standard Q-learning, which does not make any prior assumptions about the form of the utility function. In our implementation of Q-learning, the range of values the motivation can take was divided into a number of bins, and the model estimated the utility for each bin. In the simulations on each trial reinforcement *r* = 1 was received and its utility was computed using Eq. (10), which relied on the current motivation. The Q-value for the current motivation bin was updated by: Δ*Q_m_* = *α*(*U* − *Q_m_*). In Fig. 6 we compare a Q-learning approach in which the motivational state was discretised with the gradient model in our framework which does not require discretisation of the motivational state. As can be seen in in Fig. 6, both models are able to approximate the utility well. However, Q-learning takes significantly more trials to do so. Moreover, the more bins are used for the discretisation, the slower the learning occurs.

**Fig 6.**
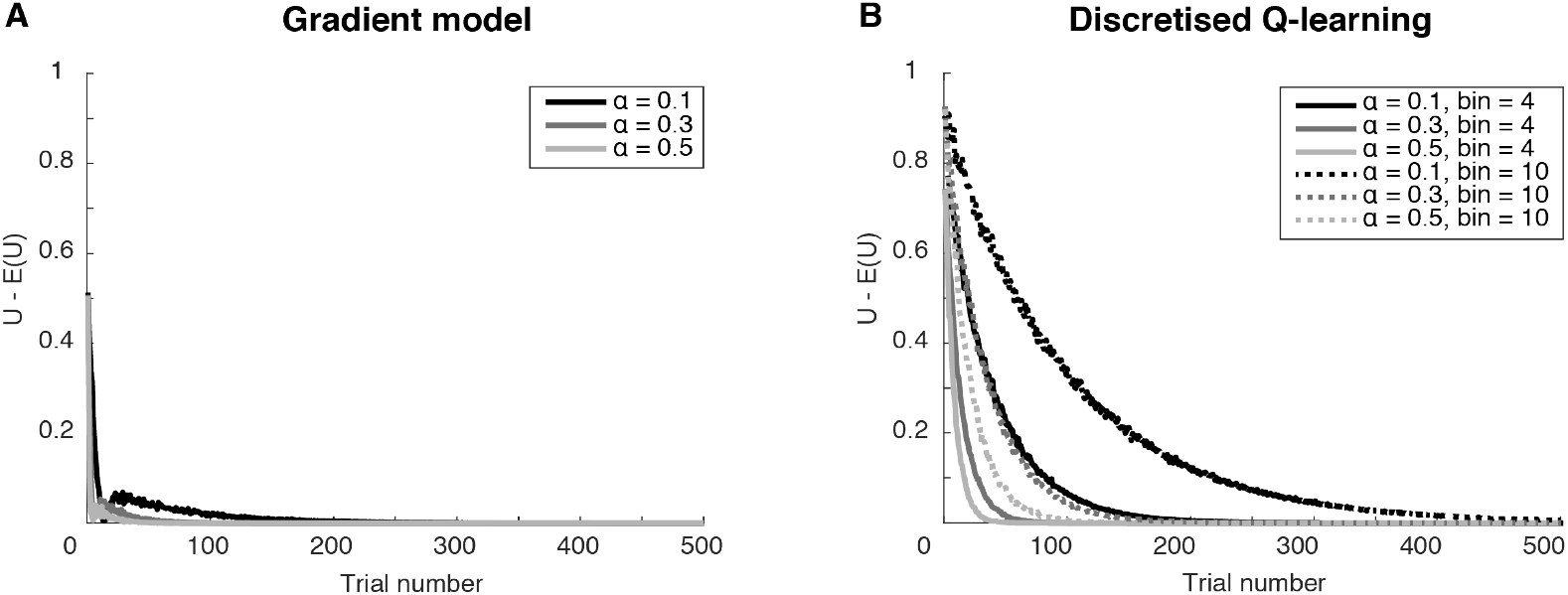
Reward prediction error as a function of learning iteration. A) gradient model using the state-dependent prediction error (Eq. (15)) and the plasticity rules described in Eq. (16) and (17). B) Discretised Q-learning model. Motivational values were randomly chosen on each trial from a uniform distribution between 0 and 2. For Q learning, motivational values were binned in either 4 or 10 bins. The *y*-axis corresponds to reward prediction error equal to the difference between the estimated and expected utility.

### Relationship to experimental data

We already demonstrated how a model employing an approximation of utility can explain data on the effect of physiological state on dopaminergic responses. In this section, we demonstrate how we can use models grounded in basal ganglia architecture to describe these dopaminergic responses and goal directed action selection in different experimental paradigms.

We first show that the new, more complex and biological relevant learning rules can also be used to explain the data by Cone et al. [25]. In these simulations, the dopaminergic teaching signal at the time of the CS took on the value of the expected utility (*T/*(1 − *D*)) and at the time of the US represented the reward prediction error described by (Eq. (15)). Simulated values of the dopaminergic teaching signal (Fig 7) show similar behaviour to the experimental data by Cone et al. [25]. Both the gradient and the payoff-cost model produce a similar dopaminergic teaching signal. This could be expected from simulations in the previous sections, which showed that both models converge to similar weights if the motivation is kept constant during training.

**Fig 7.**
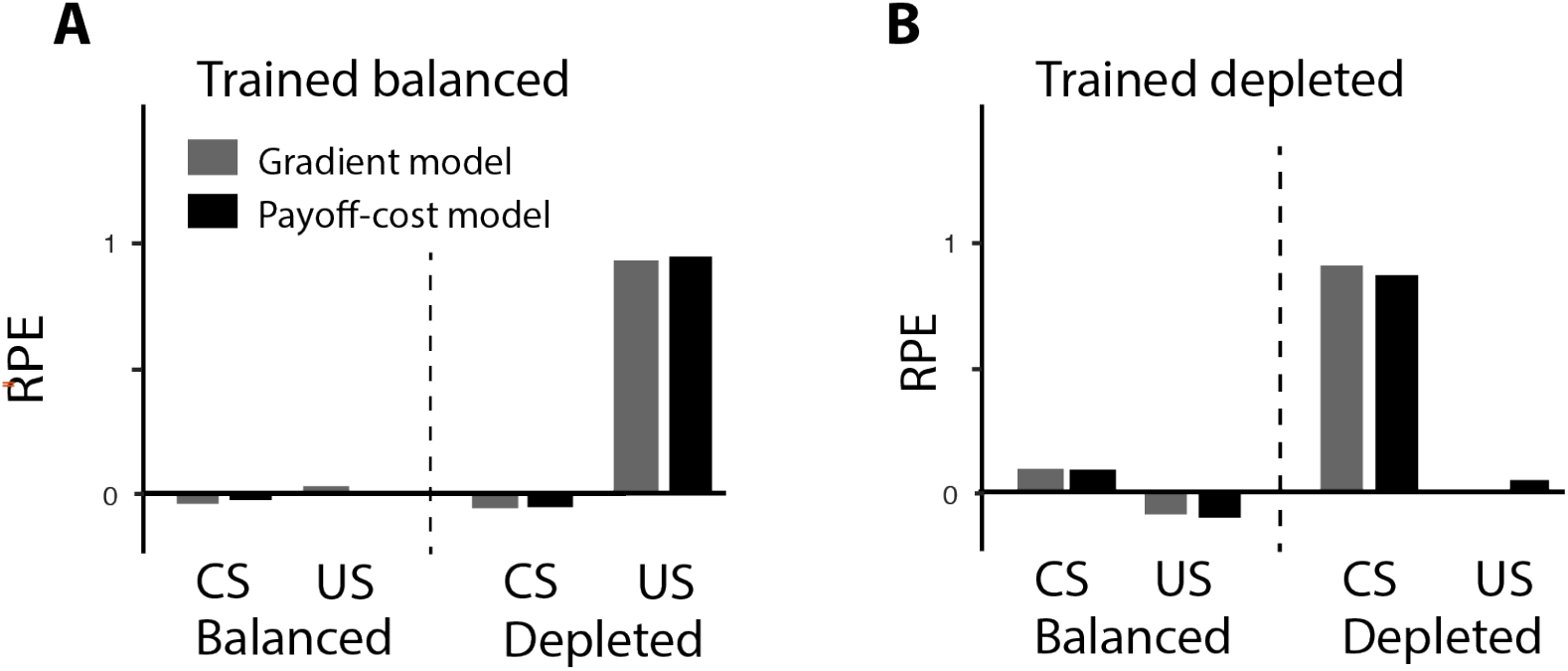
Simulated dopaminergic teaching signal in the paradigm of Cone et al. [25] according to models grounded in basal ganglia architecture. State-dependent prediction error (Eq. (15)) was used. The gradient model in grey uses the plasticity rules described in Eq. (16) and (17). payoff-cost model is depicted in black and uses the plasticity rules described in Eq. (18) and (19). Left and right panels show the data tested in the balanced state or depleted state, respectively. CS = conditioned stimulus, US = unconditioned stimulus, RPE = reward prediction error. Each simulation consisted of 50 trials and was repeated 5 times, similar to the number of animals in each group. The parameters used in the simulations were *α* = 0.1, *ϵ* = 0.8 and *λ* = 0.01.

#### Influence of physiological state on action selection

In the presented framework natural appetites, such as hunger or thirst can drive action selection into the direction of the relevant reinforcement. Generally speaking, most foods are considered appetitive even when an animal is in the near-optimal state. Nevertheless, overconsumption could have negative consequence as you can experience discomfort after eating too much. Therefore some of these negative consequences might have to be accounted for as well. As discussed above, a good example of a natural appetite that can be both appetitive and aversive dependent on the physiological state of the animal is salt appetite. Salt is considered very aversive or appetitive when the sodium physiology is balanced or depleted, respectively. Accordingly, rats reduce their intake of sodium or salt-associated instrumental responding when balanced and vice versa when depleted [26, 46]. This even occurs when animals have never experienced the deprived state before and have not had the chance to relearn the incentive value of a salt reinforcement under a high motivational state [46]. This example fits very well with the incentive salience theory which states that the learned association can be dynamically modulated by the physiological state of the animal. Modulation of incentive salience adaptively guides motivated behaviour to appropriate reinforcements.

To demonstrate that the simple utility function (Eq. (10)) proposed in this paper can account for the transition of aversive to appetitive reinforcements, and vice versa, in action selection, we use the study by Berridge and Schulkin [26]. In this study, animals learned the value of two different conditioned stimuli, one associated with salt intake (CS+) and one with fructose intake (CS-). The animals were trained when they were in a balanced state of sodium. Once the appropriate associations had been learned, the animals were tested in a sodium balanced and sodium depleted state. As can be seen in Fig 8A the intake of the CS+ was significantly increased in the sodium depleted state in comparison to the balanced state and in comparison to the CS- intake. If we assume that positive and negative consequences are encoded by the Go or Nogo pathway, respectively, the synaptic weights of these pathways will acquire positive or negative values depending on the situation. Again, the dopaminergic activation signal can control to what extent these positive and negative consequences affect the basal ganglia output as Go and Nogo neurons are modulated in an opposing manner.

**Fig 8.**
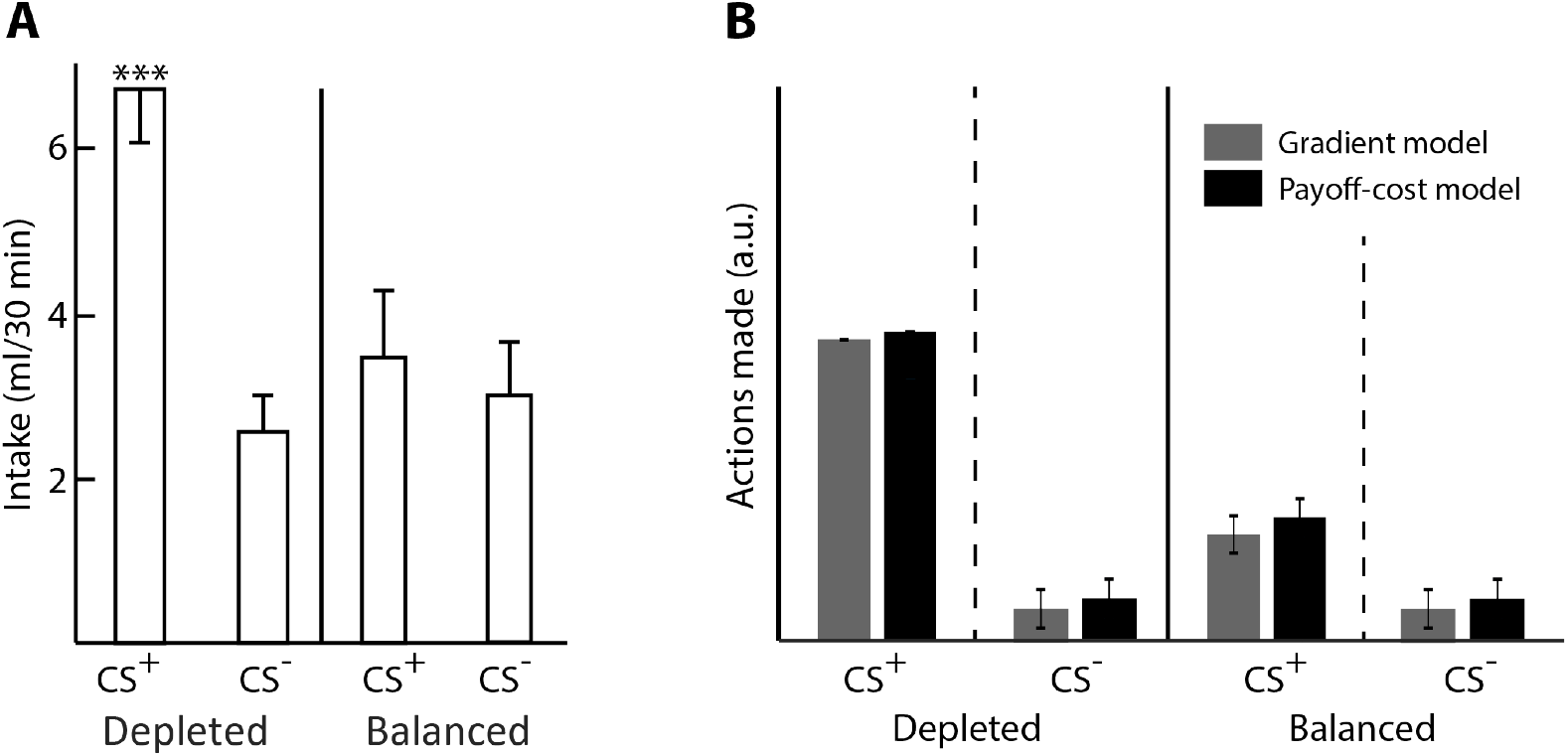
Salt appetite: experimental data by Berridge and Schulkin [26]. A) Intake of fructose (CS-) or Sodium (CS+). B) Simulated data of number of actions made using the state-dependent prediction error (Eq. 15). The gradient model in grey uses the plasticity rules described in Eq. (16) and (17). The payoff-cost model is depicted in black and uses the plasticity rules described in Eq. (18) and (19). Within each graph, the left and right halves show the responses of animals tested in depleted and balanced states, respectively. CS+ = relevant conditioned stimulus for sodium, CS- = irrelevant conditioned stimulus for fructose. Within the graph, the left and right halves show the responses of animals tested in depleted and balanced states, respectively. CS+ = relevant conditioned stimulus for sodium, CS- = irrelevant conditioned stimulus for fructose.

Once the appropriate associations between the conditioned stimuli and the outcomes are acquired, the outcomes can be dynamically modulated by the relevant state only (i.e. the level of sodium depletion). The fact that the responses to the CS- are unaffected by the physiological state of sodium suggests that salt and fructose are modulated by separate appetitive systems and that the physiological state of the animal modulates the intake proportional to the deprivational level of the animal. The phenomenon that different reward types act on different appetitive system has been also observed by other experimental studies [24].

In our simulation, we assumed that the synaptic weights for Go and Nogo neurons were learned in a near-balanced state of sodium since the animals had never experienced a sodium depleted state before. During training, the motivation was low (*m* = 0.2), resulting in low level of dopaminergic activation signal following Eq. (14). During the testing phase, the motivation for the CS+ was low (*m* = 0.2) for sodium in the near-balanced state and high (*m* = 2) for the sodium depleted state. Given that experimental data suggests that multiple appetitive systems may be involved we used separate motivational signals for the CS+ and CS-. Therefore, the motivation for the CS- were kept low (*m* = 0.1), but were non-negative, for both sodium near-balanced and sodium depleted states since fructose has no effect on the physiological state of sodium and we assumed that the animals were not deprived of other nutrients. The thalamic activity was computed using Eq. (11), and additional Gaussian noise was added to allow exploration. Actions were made when the thalamic activity was positive, otherwise no action was made. The model received a reinforcement of *r* = 0.5 for each action made and the utility was computed using Eq. (10). During training, Go and Nogo weights were updated using the update equations presented above for the different models. For the testing phase, the Go and Nogo values were kept constant based on the learned values and were not allowed to be (re-)learned. Again, the thalamic activity was computed and actions were taken when this was positive. Please note that the main difference between near-balanced and depleted states, is the level of dopaminergic activation signal. As can be seen in Fig 8B both models show dynamic scaling of the CS+ dependent on the relevant motivational state similar to the experimental data in Fig 8A.

#### State-dependent valuation

There is a number of experimental studies that have investigated the influence of physiological state at the time of learning on the preference during subsequent encounters (e.g. [27, 34, 35, 47]). In the study by Aw et al. [27], animals were trained in both a near-balanced and deprived state. One action was associated with food in the near-balanced state and another action was associated with food in the deprived state. Animals were tested in both states. In both cases, animals preferred the action associated with the deprived state during learning and the proportions of trials with these actions are above chance level (Fig 9A). These results resemble the data on dopaminergic responses (Fig 1), which also demonstrated higher response to reward-predicting stimuli (CS) that had been experienced in a depleted state. In this section we show that such preferences can be produced by the proposed models.

**Fig 9.**
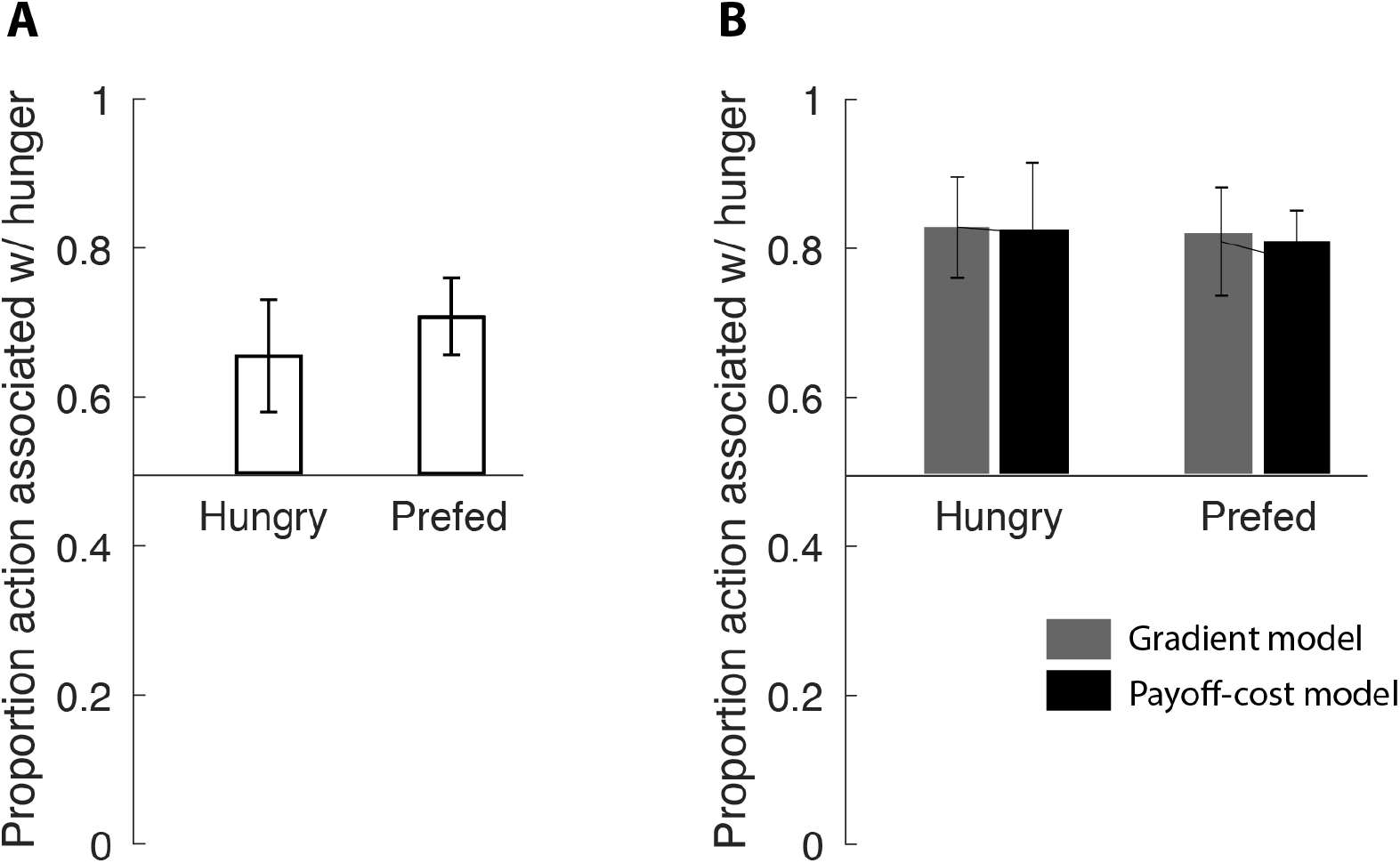
Simulation of a study by Aw et al. [27]. A) Experimental data by Aw et al. [27]. B) Simulated data using the state-dependent prediction error (Eq. 15). gradient model in grey uses the plasticty rules described in Eq. (16) and (17). payoff-cost model is depicted in black and uses the plasticity rules described in Eq. (18) and (19). Hungry and Prefed refer to the physiological state at testing. The parameters used in the simulations were *α* = 0.1, *ϵ* = 0.8 and *λ* = 0.01. Additionally a Gaussian noise with mean 0 and standard deviation 0.1 was added to the above thalamic activity.

We simulated learning of the synaptic weights of Go and Nogo neurons when the motivation was high (i.e. hungry) and when the motivation was low (i.e. prefed). In the experiment by Aw and colleagues, the training phase consisted of forced choice trials in which the reinforcement was only available in one arm of a Y-maze while the other arm is blocked. For example, the left arm was associated with a food reinforcement during hunger and the right arm was associated with a food reinforcement during the prefed condition. In the experiment, 11 animals were used, which were trained for 65 trials on average to reach the required performance. In line with this, our simulations were repeated 11 times, and in each iteration we trained the model for varying trial numbers with a mean of 65. Motivation was set to *m* = 2 for hungry and *m* = 0.2 for prefed. The dopaminergic activation signal was fixed to values that correspond to the motivation described by Eq. (14). For each correct action, the model received a reinforcement of *r* = 0.2 and the utility was calculated using the utility in Eq. (10). At the start of each simulated forced trial, the model computed the thalamic activity (using Eq. (11)) of the available action and some independent noise was added. The thalamic activity for the unavailable action was zero. The action with the highest positive thalamic activity was chosen. If the thalamic activity of all action was negative, no actions was made and the reinforcement was zero. Each time an action was made the synaptic weights of Go and Nogo neurons were updated using the state-dependent reward prediction error and the update rules described in section Models of learning. The learning rate for all of these models was set to *α* = 0.1. Once learning was completed, the synaptic weights were fixed and were not allowed to be updated anymore.

During the testing phase, both arms were available and the animals could freely choose an arm to obtain a reinforcement in. All 11 animals were tested for 24 trials. Our simulations were tested for 24 trials for both conditions and repeated 11 times using the individual learned Go and Nogo weights for the prefed and hungry condition. Again, the model computed the thalamic activity for both options simultaneously (in parallel) plus some independent noise. The action with the highest thalamic activity was chosen. The proportion of actions associated with the hungry option are depicted in Fig 9B. This experiment was simulated for both physiological states during testing phase. The proportion of actions for the arm associated with hunger were calculated for both states. Both the experimental and simulated data show that the animals chose the action associated with the hungry state more often, regardless of the current state.

To gain some intuition for why the models preferred the option that was associated with hungry state during training, let us look back at the simulations presented in Fig 5B-D. They show that when the models were trained with a fixed motivation, Go weights took higher values when the motivation was high during training, and Nogo weights were larger when motivation was low. Analogously, in the simulations of the study of Aw and colleagues, the Go weights took larger values for the option associated with hunger during training (Fig 10), and this option was therefore preferred during testing. These biases arise in the models when they are unable to experience multiple levels of motivation during training. Only after training in multiple physiological states the utility can be flexibly estimated in different physiological states.

**Fig 10.**
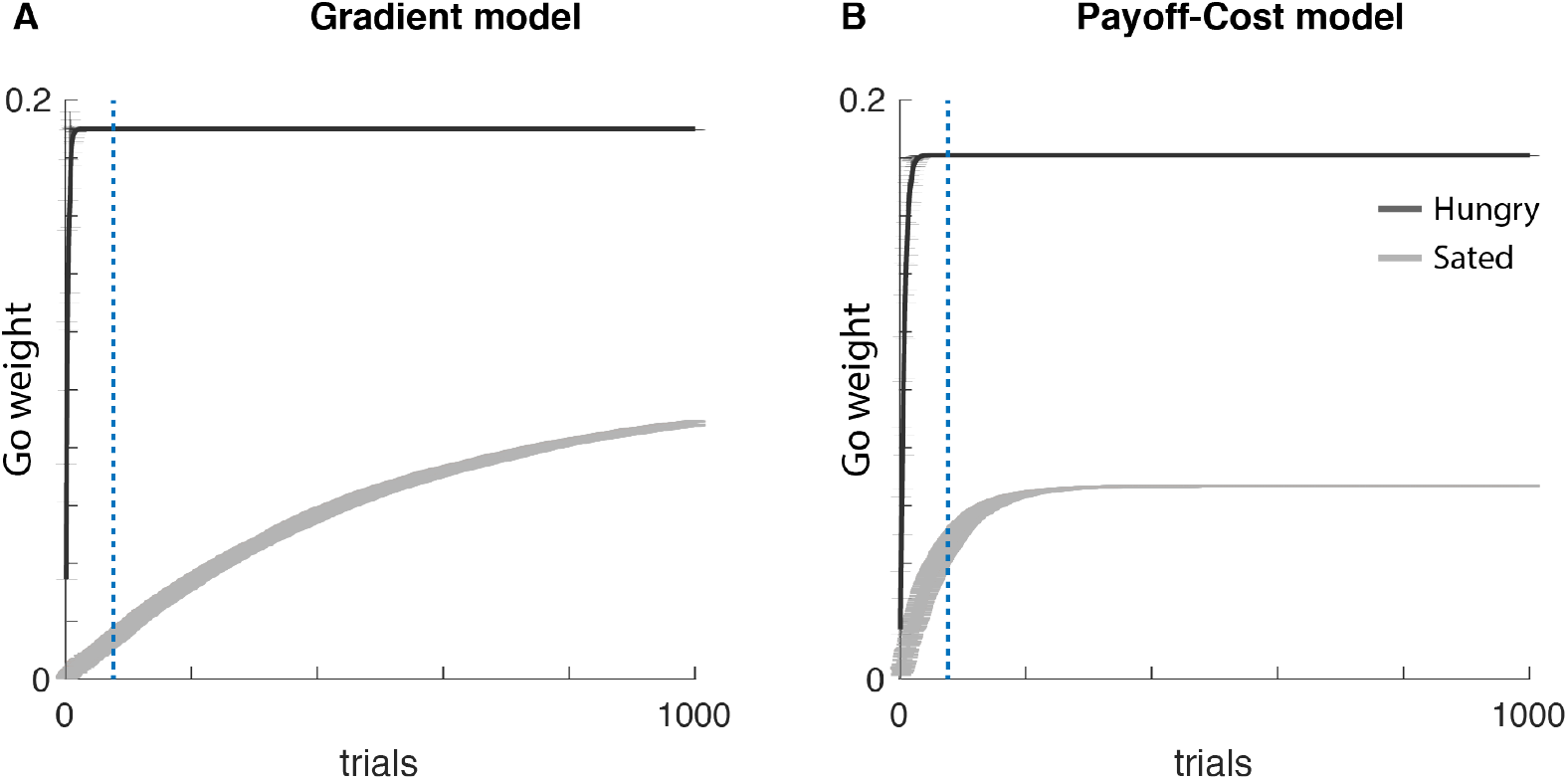
Learning Go weights as function of physiological state. Learning at high motivation is depicted in black and learning at low motivation is depicted in grey. Number of trials used for the simulation was 1000. Go weights were initialised at zero. The parameters used in the simulations were *α* = 0.1, *ϵ* = 0.8 and *λ* = 0.01. Additionally a Gaussian noise with mean 0 and standard deviation 0. 1 was added to the above thalamic activity.

## Discussion

In this paper, we have presented a novel framework for action selection under motivational control of internal physiological factors. The framework is biologically grounded and brings together models of direct and indirect pathways of the basal ganglia with the incentive learning theory. We proposed two models that learn about positive and negative aspects of actions utilising a prediction error that is influenced by the current physiological state. In this section, we will discuss the experimental predictions, the relationship to experimental data and other computational models and other implications.

### Experimental predictions

In this section, we outline the predictions the models make. The neural implementation of the framework assumes that Nogo neurons prevent selecting actions with large reinforcements when the motivation is low. Thus it predicts that pharmacological manipulations of striatal Nogo neurons through D2 agonist (or antagonist) should increase (or decrease) the animal’s tendency to consume large portions of food or other reinforcers to a larger extent when it is close to satiation, than when it is deprived.

The neural implementation of the framework also assumes that the activity in Go and Nogo pathways is modulated by the dopaminergic activation signal, which depends on motivation. This assumption could be tested by recording activity of Go and Nogo neurons, for example using photometry, while an animal decides whether to consume a reinforcement. The framework predicts that deprivation should scale up responses of Go neurons, and scale down the response of Nogo neurons.

As showed in Fig. 5A, the framework predicts that the synaptic weights of Go and Nogo neurons converge to different values depending on the reinforcement magnitude. These predictions can be tested in an experiment equivalent to the simulation in Fig. 5A in which mice learn that different cues predict different reinforcement sizes, and experience each cue in variety of motivational states. The weights of the Go and Nogo neurons are likely to be reflected by their neural activity (e.g. measured with photometry) while an animal evaluates a cue at baseline motivation level. We expect both populations to have higher activity for cues predicting higher reinforcements, and additionally, the gradient model predicts that the Go and NoGo neurons should scale their activity with reinforcement magnitude linearly and quadratically, respectively.

### Relationship to experimental data and implications

The proposed framework can account for decision making and learning as a function of physiological state, as shown by the simulations of the data by Cone et al., Berridge and Schulkin and Aw et al. More specifically, we proposed that learning occurs based on the difference between the utility and expected utility of an action. This is in line with results from a study in monkeys that also suggested that dopaminergic responses reflects a difference in utility of obtained reward and expected utility [30]. That study focused on a complementary aspect of subjective valuation of reward, namely that the utility of different volumes of reward is not equal to the objective volume, but rather to its nonlinear function. In this paper we additionally point out that the utility of rewards depends on the physiological state in which they are received.

Furthermore, we know from literature that low levels of dopamine, as seen in Parkinson’s disease patients, drive learning from errors, whereas normal/high levels dopamine emphasise positive consequences [3, 5, 48]. In our simulations, we observe this as well: a low dopaminergic activation signal emphasises the negative consequences of actions encoded in the synaptic weights of Nogo neurons, whereas a high dopaminergic activation signal emphasises the positive consequences of actions encoded in the synaptic weights of Go neurons. However, there are a couple considerations that have to be made with respect to the dopaminergic signal in our simulations. First, we assume that striatal neurons can read out both motivational and teaching signals encoded by dopaminergic neurons [21]. In our theory, we describe two roles of dopamine neurons, namely activation and teaching signal, however, we do not provide a solution to how these different signals are accessed. The function of dopamine neurons has been under a current debate and its complexity is not well understood [22]. We will leave the details of the mechanisms by which they can be distinguished to future work. We assume that the models, particularly the gradient model, has access to multiple dopaminergic signals simultaneously. Although we recognise that this is a simplified concept of what might be happening in the brain, it still provides us with new insights in how these different functions affect aspects of decision making. Further research is necessary to describe the complexity of dopamine neurons in decision making.

Secondly, in this paper we have focused primarily on one dimension, namely nutrient deprivation. However, experimental data suggests that reinforcements are scaled selectively by their physiological needs [24]. A nutrient specific deprivation alters goal-directed behaviour towards the relevant reinforcement, but not the irrelevant one. In contrast, other physiological factors, such as fatigue, may scale only the negative, but not the positive consequences. This hypothesis is supported by data showing that muscular fatigue alters dopamine levels [49]. Together this suggests that the utility of an action is most likely the sum of all the positive and negative consequences with respect to their physiological needs or other external factors. Therefore, extending the current theory to multiple dimensions is an important direction of future work. In such an extended model, an action which changes the state of multiple physiological dimensions, e.g. hunger, thirst and fatigue, would need to be represented by multiple populations of Go and Nogo neurons. In this example, the value of food and drink reinforcement *r_f_* and *r_d_* would need to be represented by separate populations of Go and Nogo neurons modulated by different populations of dopaminergic neurons encoding information about hunger and thirst, while the effort would need to be encoded by a population of Nogo neurons modulated by a fatigue signal. It would be interesting to investigate if the required number of neurons could be somehow reduced by grouping terms in the utility function scaled in a similar way (e.g. 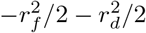, which are not scaled by any factor in this example). Future experimental work on the diversity of how individual dopaminergic neurons are modulated by different physiological states would be very valuable in constraining such an extended model.

Although the current study does not focus on risk preference, there is some evidence for the existence of a link between risk preference and physiological state [32, 50]. Particularly, in the payoff-cost model the dopaminergic activation signal controls the tendency to take risky actions [42], thereby predicting that motivational states such as hunger can increase risk seeking behaviour. The above mentioned studies show that changes in metabolic state systematically alter economic decision making for water, food and money and correlate with hormone levels that indicate the current nutrient reserve. Individuals became more risk-averse when sated whereas people became more risk-seeking when food deprived.

The current study also provides insights into the mechanistic underpinnings of overeating and obesity. Imaging studies using positron emission tomography showed an important involvement of dopamine in normal and pathological food intake in humans. In comparison to healthy controls, pathologically obese subjects show reduced availability of striatal D2 receptors that were inversely associated with the weight of the subject [51–53]. Our theory suggests that the ability to restrain from taking actions and learn from negative consequences of actions such as overeating may be diminished when D2 receptors are activated to a lesser extent. The involvement of the DA system in reward and reinforcement suggests that low engagement of Nogo neurons in obese subjects predisposes them to excessive use of food.

### Relationship to other computational models

The proposed framework builds on or is related with several other theories. For example, Keramati and Gutkin [28] developed a theory that also extended the reinforcement learning theory to incorporate physiological state. They defined a ‘homeostatic space’ as a multidimensional metric space in which each dimension represents physiologically-regulated variable. At each time point the physiological state of an animal can be represented as a point in this space. They also define motivation (to which they refer to as ‘drive’) as the distance between the current internal state and the desired setpoint. We extended this theory to include how the brain computes the modulation of learned values by physiology.

In the motivation for the existence of the desired physiological state, Keramati and Gutkin [28] referred to active inference theory [54]. Our framework also shares a conceptual similarity with this theory, in that both action selection and learning can be viewed as the minimisation of surprise. To make this link clearer, let us provide a probabilistic interpretation for action selection and learning processes in our framework. This interpretation is inspired by a model of homeostatic control [55]. It assumes that the animal has a prior expectation *P*(*S*) of what the physiological state *S* should be, which is encoded by a normal distribution with mean equal to the desired state *S*\*. That model assumes that animals have an estimate of their current bodily state *S* (interoception). It proposes that animals avoid states *S* that are unlikely according to the prior distribution with mean *S*\* (thus they minimise their “interoceptive surprise”), and they wish to find themselves in the states *S* with high prior probability *P*(*S*). Following these assumptions, we can define the desirability of the state as *Y*(*S*) = ln*P*(*S*). If we assume for simplicity that *P*(*S*) has unit variance and ignore an additive constant, we obtain our definition of a desirability of a state in Eq. (2). In our framework, actions are chosen to minimise the surprise of ending up in a new physiological state. The closer this state is to the desired state the more likely it will be and the smaller the surprise. Furthermore, motivation itself could be viewed as an error in the prediction of the physiological state.

Similar to action selection, animals update the parameters of their internal model (e.g. *V*, *G*, *N*) during learning in order to be less surprised by the outcome of the chosen action. To describe this more formally, let us assume that the animal expects the utility to be normally distributed with mean *Û* and variance 1 (for simplicity). Furthermore, assume that during learning the animal minimises the surprise about the observed utility of action *U*. Therefore, we can define the negative of this surprise as *F* = ln*P*(*U*). This objective function is equal (ignoring a constants) to our objective function defined in Eq. (8). Thus in summary, similar to the active inference framework, both action selection and learning could be viewed as processes of minimising prediction errors.

The dopaminergic activation signal is often associated with an increase in the vigour of actions [10]. In the study by Niv et al. [10] same assumption is held that the utility of the reinforcement is dependent on the deprivational level, however, they do not provide a mechanism for how these utilities are computed and are therefore set them arbitrarily. Moreover, they rely on average reward reinforcement learning techniques which reveal an optimal policy that leads to an average reward rate per time unit. Following this line of thinking, actions with higher utility (i.e. actions taken in a deprived state) cause higher response rates as the opportunity cost of time increases. Although our model does not describe vigour or response times, it could be related to these output statistics thanks to recent work investigating the relationship between activity of a basal ganglia model and the parameters of a diffusion model of response times in a two alternative choice task [56]. This study showed that a drift parameter of a diffusion model is related to the difference in the activation of Go neurons selective for the two options, while the threshold is related to the total activity of Nogo neurons. In our framework motivation scales linearly with Go neurons for both options; it enhances the difference in their activity. Based on the data by Dunovan et al. [56] motivation is expected to increase the drift rate and reduce the threshold leading to faster responding.

There are many computational models developed for action selection in either very abstract or more biological relevant ways. One of the leading models in describing how dopamine controls the competition between the Go and Nogo pathways during action selection is the Opponent Actor Learning (OpAL) model. This model hypothesises that the Go and Nogo neurons encode the positive and negative consequences of actions respectively [41] and high dopamine levels excite Go neurons and low levels of dopamine releases the inhibition of Nogo neurons. Moreover, existing neurocomputational theories describe how experience modifies striatal plasticity and excitability of the Go and Nogo neurons as a function of reward prediction errors [13, 41, 42, 44]. In line with these studies, we assumed that the Go neurons encode positive consequences and Nogo neurons encode negative consequences and that dopaminergic activation signal controls the balance between these neurons. We extended these concepts by combining it with incentive salience theory.

Our framework considers for simplicity that all physiological dimensions (e.g. hunger, salt level, body temperature) have unique values with a maximum desirability, and the desirability is lower for both smaller and higher values along the physiological dimension. Although this assumption is realistic for some dimensions, it may not be realistic for the resources animals store outside their bodies. Indeed, according to the classical economic utility theory, humans always wish to have more monetary resources. Although our framework also assumes diminishing utility of larger reinforcements, it differs from the classical theory in that the utility is a non-monotonic function of reinforcement. We also assume that all the resources are directly consumed, which does not allow for the scenario of storing resources. It would be interesting to extend presented model to include dimensions without finite value maximising desirability.

Up to date, there is only a few neurocomputational studies on incentive salience (e.g. [37, 57]). In these studies two mechanisms are proposed on how the physiological state may influences reinforcement evaluation [37]. In the study by Zhang et al., [37], two mechanisms are proposed through which a physiological state, *κ*, modulates reinforcement, *r*: one mechanism is additive (*r* + log(*κ*)) and the other is multiplicative (*rκ*). Appetitive reinforcements are considered positive and aversive reinforcements are considered negative. The physiological state of the animals is always non-negative, *κ* ∈ [0, ∞). The multiplicative mechanism can perfectly account for either positive or negative reinforcements, but struggles to explain a phenomenon such as salt appetite where aversive reinfrocements can become appetitive depending on the physiological state of the animal. This is where the additive mechanism comes in. This mechanism is able to change the polarity of the reinforcement value without changing the sign of the reinforcement. We build on this line of thinking and propose only one mechanism for incentive salience that can account for positive and negative reinforcements without the need to change the sign of the reinforcement. Our utility theory accounts for positive and negative consequences of actions in a state-dependent manner. Moreover, even when the model learns reinforcement values in a non-depleted state and has never experienced a depleted state before, the model is able to behave appropriately. In conclusion, our modelling framework maps learning of incentive salience onto the basal ganglia circuitry, a circuitry proven to play an important role in action selection. We used key concepts from both lines of theoretical work to develop a framework that is biologically relevant and describes action selection and learning in a state-dependent manner.

## Acknowledgments

We thank Moritz Möller for his feedback on the manuscript.

